# Decreased efficacy of a COVID-19 vaccine due to mutations present in early SARS-CoV-2 variants of concern

**DOI:** 10.1101/2023.06.27.546764

**Authors:** Payton A.-B. Weidenbacher, Natalia Friedland, Mrinmoy Sanyal, Mary Kate Morris, Jonathan Do, Carl Hanson, Peter S. Kim

## Abstract

With the SARS-CoV-2 virus still circulating and evolving, there remains an outstanding question if variant-specific vaccines represent the optimal path forward, or if other strategies might be more efficacious towards providing broad protection against emerging variants. Here, we examine the efficacy of strain-specific variants of our previously reported, pan-sarbecovirus vaccine candidate, DCFHP-alum, a ferritin nanoparticle functionalized with an engineered form of the SARS-CoV-2 spike protein. In non-human primates, DCFHP-alum elicits neutralizing antibodies against all known VOCs that have emerged to date and SARS-CoV-1. During development of the DCFHP antigen, we investigated the incorporation of strain-specific mutations from the major VOCs that had emerged to date: D614G, Epsilon, Alpha, Beta, and Gamma. Here, we report the biochemical and immunological characterizations that led us to choose the ancestral Wuhan-1 sequence as the basis for the final DCFHP antigen design. Specifically, we show by size exclusion chromatography and differential scanning fluorimetry that mutations in the VOCs adversely alter the antigen’s structure and stability. More importantly, we determined that DCFHP without strain-specific mutations elicits the most robust, cross-reactive response in both pseudovirus and live virus neutralization assays. Our data suggest potential limitations to the variant-chasing approach in the development of protein nanoparticle vaccines, but also have implications for other approaches including mRNA-based vaccines.

## INTRODUCTION

DCFHP-alum is a SARS-CoV-2 spike-functionalized, ferritin nanoparticle-based vaccine candidate that is formulated with aluminum hydroxide (alum) as the sole adjuvant^1^. The antigen component of the vaccine contains a 70 amino acid C-terminally truncated spike ectodomain (∆C70), containing the HexaPro^2^ mutations that stabilize the prefusion state (DCFHP; Delta-C70-Ferritin-HexaPro). Previously, we demonstrated that DCFHP-alum, when administered with a prime-boost regimen, elicits potent neutralizing antisera in non-human primates (NHPs) against all tested SARS-CoV-2 variants of concern (VOCs), including Omicron strains, as well as against SARS-CoV-1. The neutralizing antibody response was durable, and all NHPs maintained neutralizing antisera for over 250 days. To explicitly test the potential of DCFHP-alum as an annual vaccine, NHPs were given a booster ∼one year after the initial immunization; all animals showed a strong anamnestic neutralizing immune response against SARS-CoV-2 VOCs and SARS-CoV-1. Our results suggest that DCFHP-alum has potential to be used as a once-yearly (or less frequent) booster vaccine, and as a primary vaccine for pediatric use, including in infants^1,3^.

While DCFHP is based on the Wuhan-1 sequence, we wished to investigate whether SARS-CoV-2 spike protein variants would be more efficacious. We therefore investigated different variant forms of DCFHP, containing amino acid substitutions in the spike protein encoded in different SARS-CoV-2 VOCs: (i) D614G, a mutation that rapidly dominated the global population in 2020; (ii) B.1.429, first identified in California and known as the Epsilon variant; (iii) B.1.1.7, first identified in the United Kingdom and known as the Alpha variant; (iv) P1, first identified in Brazil and known as the Gamma variant; and (v) B.1.351, first identified in South Africa and known as the Beta variant (Figure 1)^4^. This work was conducted prior to the waves of infection caused by the Delta and Omicron VOCs^4^. To investigate the relative efficacy of these different DCFHP-variants, we cloned, expressed, and purified each of these variant vaccine candidates contemporaneously with the original DCFHP, to profile their biochemical and immunological properties.

**Figure 1.**
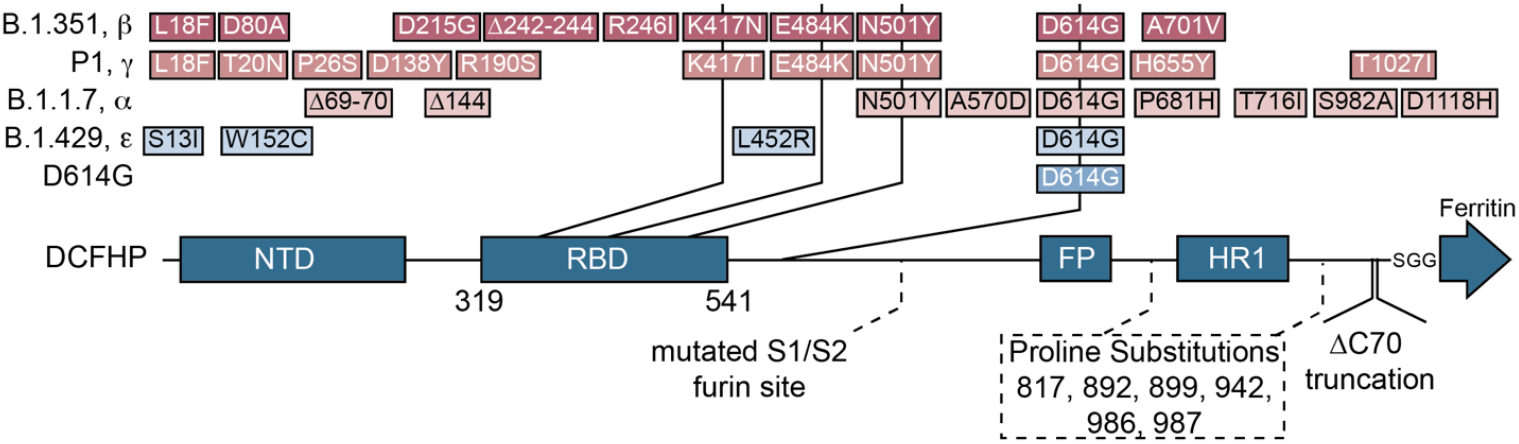
Schematic to indicate variant forms of DCFHP. The original DCFHP sequence with amino acid substitutions from the Wuhan-1 spike ectodomain is indicated on the bottom, with residue numbers and mutations labeled. Dashed box indicates residues for HexaPro mutations. Mutations in the VOCs indicated above, with common mutations joined by vertical lines. Most evolutionarily distant variants shown at top.

Here we report the expression, purification, thermal stability, and immunogenicity profiles of these vaccine candidates. While the expression of all vaccine candidates in mammalian cells was similar, size-exclusion chromatography (SEC) indicated differences in the homogeneity of the purified nanoparticles. In addition, different vaccine candidates exhibited substantially different thermal unfolding profiles, as determined by differential scanning fluorimetry (DSF). Strikingly, immunization with a vaccine candidate based on a particular VOC did not always elicit the highest neutralization titers against that VOC. Indeed, overall, immunization with the original DCFHP, based on the Wuhan-1 strain, elicited the most cross-reactive neutralizing titers. Our results highlight that some variant-specific vaccine candidates may be suboptimal future vaccines.

## RESULTS

### Vaccine design

Nanoparticle vaccines optimally facilitate increased uptake by dendritic cells and promote B-cell receptor clustering, leading to enhanced immunogenicity compared to isolated subunit vaccines^5,6^. We previously produced our vaccine candidate DCFHP by engineering the Wuhan-1 strain of SARS-CoV-2 spike protein and displaying it on a ferritin nanoparticle scaffold^1,3,7^. DCFHP presents the engineered spike protein at the three-fold axes of symmetry on the ferritin nanoparticle, enabling assembly of eight spike trimers from the constituent 24 monomers. Our design includes the removal of the 70 amino acids at the C-terminus of the ectodomain, an immunodominant region of the spike protein known to elicit non-neutralizing antibodies^8-10^. The design also includes six prefusion-stabilizing proline substitutions found in HexaPro, as well as mutations within the furin cleavage site that prevent cleavage between the S1 and S2 domains^2,11,12^. The proline modifications stabilize the spike protein in the native, prefusion conformation, and the C-terminal truncation also decreases the flexibility of the spike protein to optimally orient the RBD domain radially from the ferritin core. We generated five DCFHP-variants, in the DCFHP background, by incorporating mutations from the VOCs D614G, Epsilon, Alpha, Gamma, and Beta (Figure 1), to produce DCFHP-G, DCFHP-γ, DCFHP-α, DCFHP-γ, and DCFHP-β, respectively (Figure 1).

### Expression and purification of DCFHP and DCFHP-variant vaccine candidates

We expressed and purified DCFHP-variants utilizing the same protocol as for DCFHP^1^ and in each case collected the same fractions from the final SEC purification corresponding to the front end of the nanoparticle peak. All six nanoparticles had comparable yields (Figure 2, left) and showed similar SDS-PAGE profiles (Figure 2, right). However, while DCFHP can be purified as a homogeneous species and runs as a single peak at 3.4 MDa, as determined by size exclusion chromatography and multi-angle light scattering analysis (SEC-MALS)^1^, the SEC profiles for the DCFHP-variants were not all uniformly homogenous (Figure 3). DCFHP-variants DCFHP-β and DCFHP-γ ran as less homogenous nanoparticles, with a later elution, and with larger shoulders, indicating decreased particle integrity of these DCFHP-variants (Figure 3).

**Figure 2.**
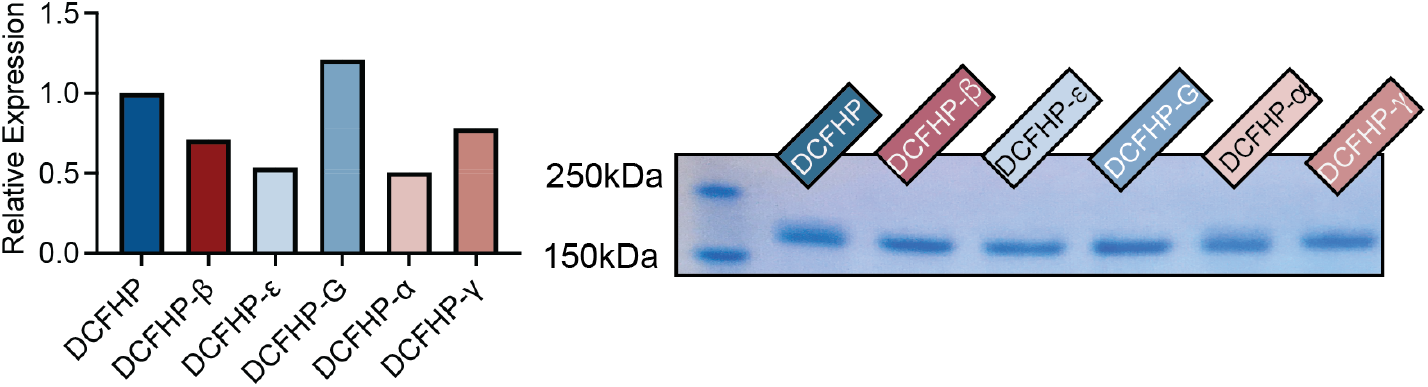
Nanoparticles were produced at a similar yield and nanoparticle monomers are of similar molecular weight. Left, relative expression of protein nanoparticles as determined by quantitation of SDS-PAGE bands from expression cultures shows that all nanoparticles are produced at a similar yield. Right, SDS-page analysis of DCFHP-variants shows that proteins are pure and with similar molecular weights. Color coding of variants matches that in Figure 1.

**Figure 3.**
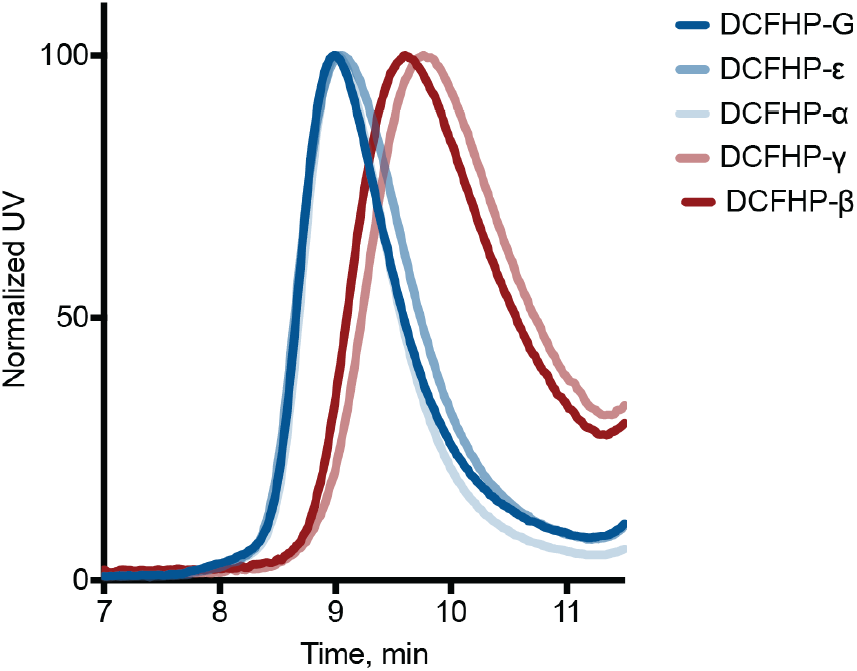
Altered nanoparticle structure of DCFHP variants. SEC was conducted with purified DCFHP-variants using an SRT-1000 column. Elution at approx. 9 min is consistent with purified DCFHP. The DCFHP-β and DCFHP-γ variants elute at a higher volume than where DCFHP and the other variants elute.

### Antigenicity and stability of DCFHP and DCFHP-variants

Consistent with literature reports, we found using biolayer interferometry (BLI) that the DCFHP-β and DCFHP-γ variants bound with higher affinity to the SARS-CoV-2 receptor ACE2 (in the form of Fc-ACE2) compared to DCFHP^13^ (Figure 4). We next tested the binding of the variants to conformation-specific monoclonal antibodies (mAbs): a class 2 RDB-binding antibody^14^ CoVA2-15^15^, a class 4 RBD-binding antibody^16^ CR3022^17^, and the a class 1 RBD-binding mAb CB6^18^ (the parent of etesevimab)^19,20^. The DCFHP-variants show the anticipated antigenicity, where binding of CB6 is substantially reduced against DCFHP-β, consistent with mutations in the RBD, CR3022 showed only limited changes in binding^21^, and the class 2 mAb, CoVA2-15, shows decreased binding to E484K containing variants (DCFHP-β and DCFHP-γ)^14,22^. Collectively, these data demonstrate that the RBD components of the DCFHP-variants display expected antigenic properties (Figure 4).

**Figure 4.**
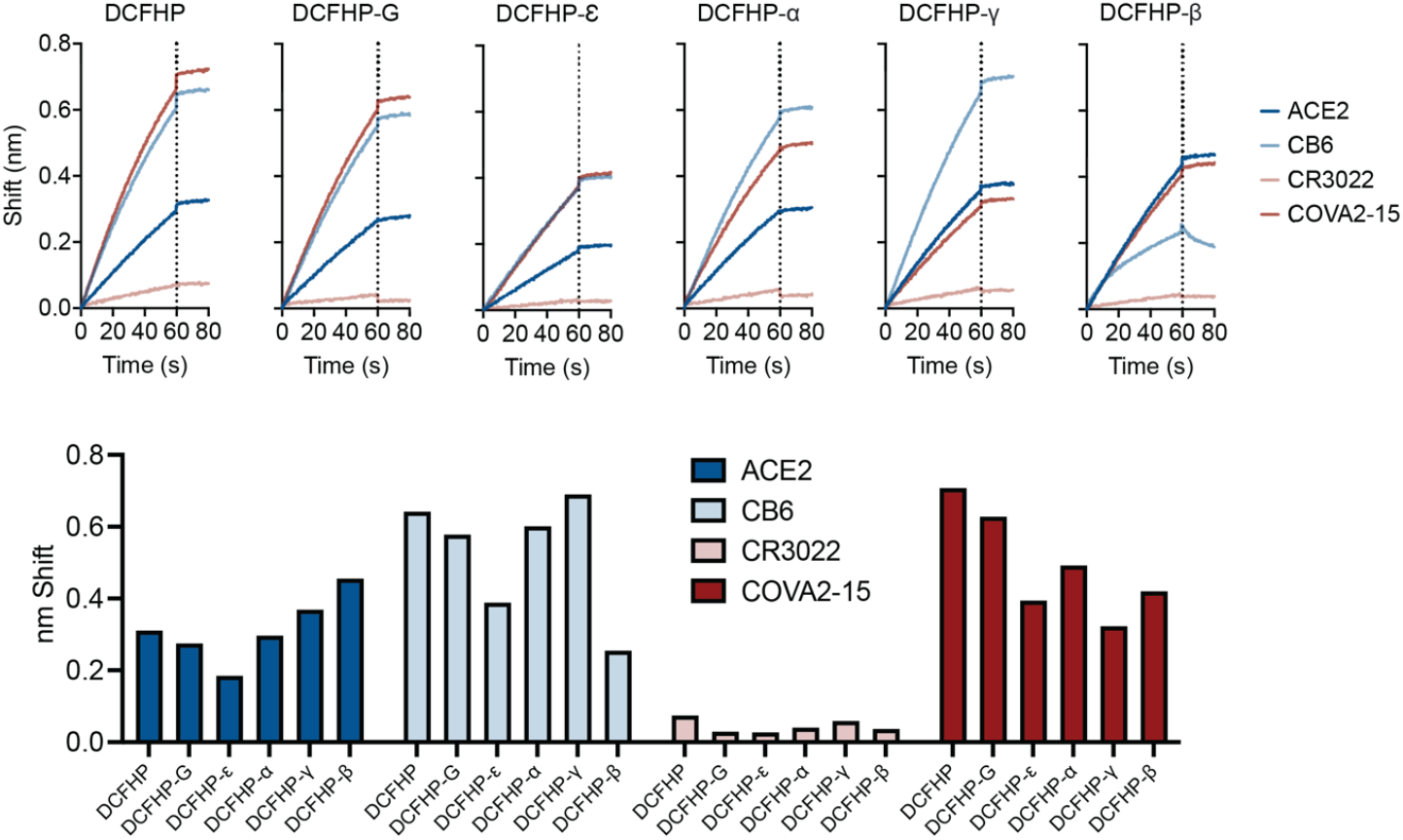
Binding of DCFHP and its variants by conformation-specific and RBD-binding mAbs correspond to their expected antigenicity. Biolayer interferometry (BLI) analysis of DCFHP and DCFHP-variants show binding to Fc-ACE2 and mAbs to differing extents. (Top) BLI shifts in nm indicated on left, time since start of association shown on bottom, dotted line indicates the beginning of the dissociation step. (Bottom) BLI shifts at the end of the association step for individual mAbs or Fc-ACE2 for different DCFHP-variants.

We previously reported an analysis of DCFHP stability using differential scanning fluorimetry (DSF), which showed two major thermal melting transitions (Tms), reminiscent of the thermal transitions reported for the stabilized HexaPro trimer^2^, providing evidence that the spike component of DCFHP is retained in a stable and native structure on the ferritin nanoparticle scaffold^1^. We assessed the melting profiles of the DCFHP-variants here and found several different profiles (Figure 5). Strikingly, some candidates yielded melting curves with altered profiles at both Tms, demonstrating that even a few mutations throughout the spike protein can impact overall protein stability. We speculate that this altered stability may contribute to the increased infectivity of some VOCs^23^. We also proceeded to test whether this altered stability affects the immunogenicity of the VOCs and/or the specificity of the immune response.

**Figure 5.**
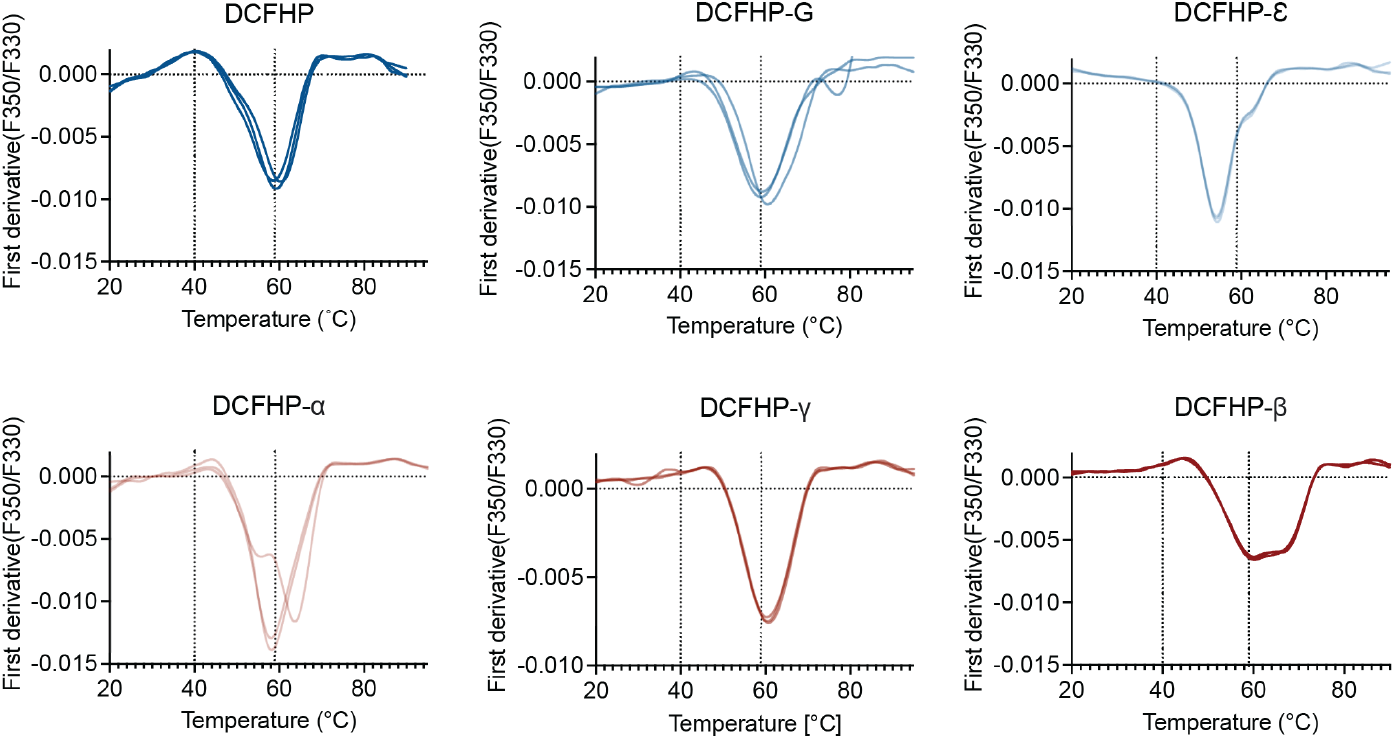
Variations in the thermal melting profiles of DCFHP variant proteins. Thermal melting data from DCFHP and variant proteins were monitored using differential scanning fluorimetry. Melting temperatures were determined by peaks of the first derivatives of the ratio of F_350_/F_330_. Altered melting profiles indicate changes in protein thermal/conformational stability. Three replicates are shown as overlaid curves. Color coding of variants matches that in Figure 1.

### DCFHP based on Wuhan-1 elicits a stronger and more broadly neutralizing immune response than more recent VOCs

To investigate the relative immunogenicity of the five DCFHP-variants, we immunized mice with the original DCFHP antigen or each DCFHP-variant, adjuvanted with alum and CpG. At day 21, following a single-dose prime, pooled antisera from each group of immunized mice were analyzed using ELISA for their binding titers to different recombinantly expressed variant spike-protein trimers. If immunization with DCFHP-variants elicit antisera that bind preferably to the matched antigen, we would expect to observe a “diagonal” in the heat map comparing binding to the variant spike-protein trimers and the DCFHP-variant used for vaccination. However, the data were more complex. Besides not observing the diagonal profile of binding, we also found that animals immunized with DCFHP-ε showed the highest overall titers against the broadest range of spike variants in this ELISA format, while those immunized with DCFHP-β showed the weakest binding to spike-protein trimers (Figure 6).

**Figure 6.**
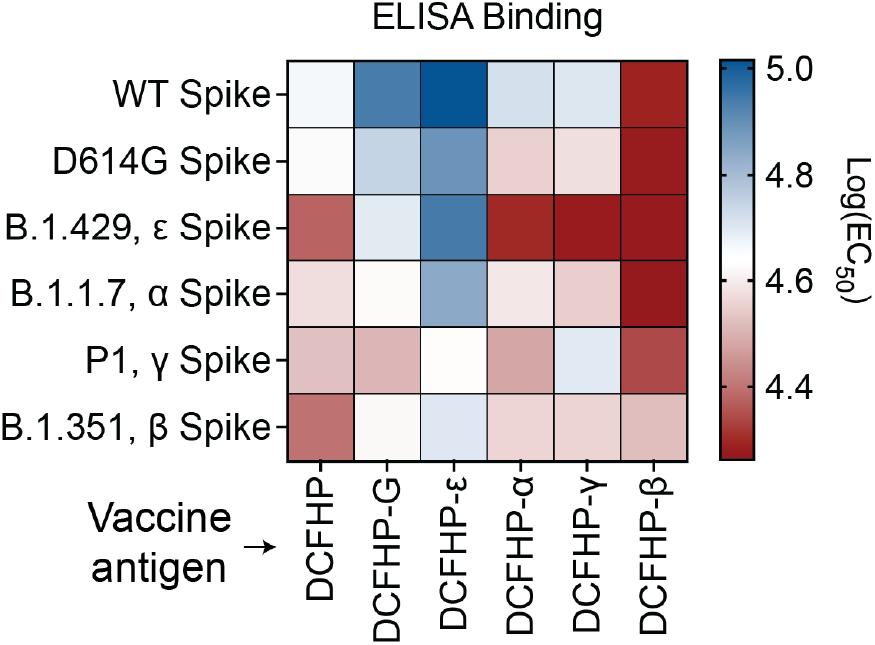
Antisera elicited by DCFHP and its variants show unpredictable binding to matched antigens. Pooled sera from mice 21 days following initial immunization with DCFHP-variants (x-axis) were tested for binding to spike protein-trimers of variant viruses (y-axis), using ELISA, and the immune response profiles plotted as a heatmap. Blue color indicates better binding of pooled anti-sera (scale shown on right). n = 5 animals included in each pool.

Next, we tested the pooled antisera from DCFHP and DCFHP-variant vaccinations for neutralization of viruses pseudotyped with the panel of VOCs. Similar to the results of the binding data in Figure 6, we did not observe a “diagonal” trend in the heat map for neutralization of pseudotyped VOCs based on the DCFHP-variant used for vaccination (Figure 7). Unlike binding, however, DCFHP-ε did not show optimal neutralizing activity. Instead, the DCHFP-variant antisera, in most cases, underperformed that from the original DCFHP vaccine. For example, antisera elicited by DCFHP-β and DCFHP-γ neutralize their corresponding viruses more weakly than antisera elicited by DCFHP did against those viruses (Figure 7). Indeed, except for neutralization of D614G, mice immunized with DCFHP showed equivalent or improved titers against all VOCs than mice immunized with the matched DCFHP-variants (either on day 21 with only a prime, or on day 28 with a prime and a boost). Testing of individual animal serum samples against the Wuhan-1 strain confirmed that the trends observed with the pooled sera were similarly reflected with individual animals, and not due to potent outlier animal(s) in the DCFHP-immunized group (Figure 7, top).

**Figure 7.**
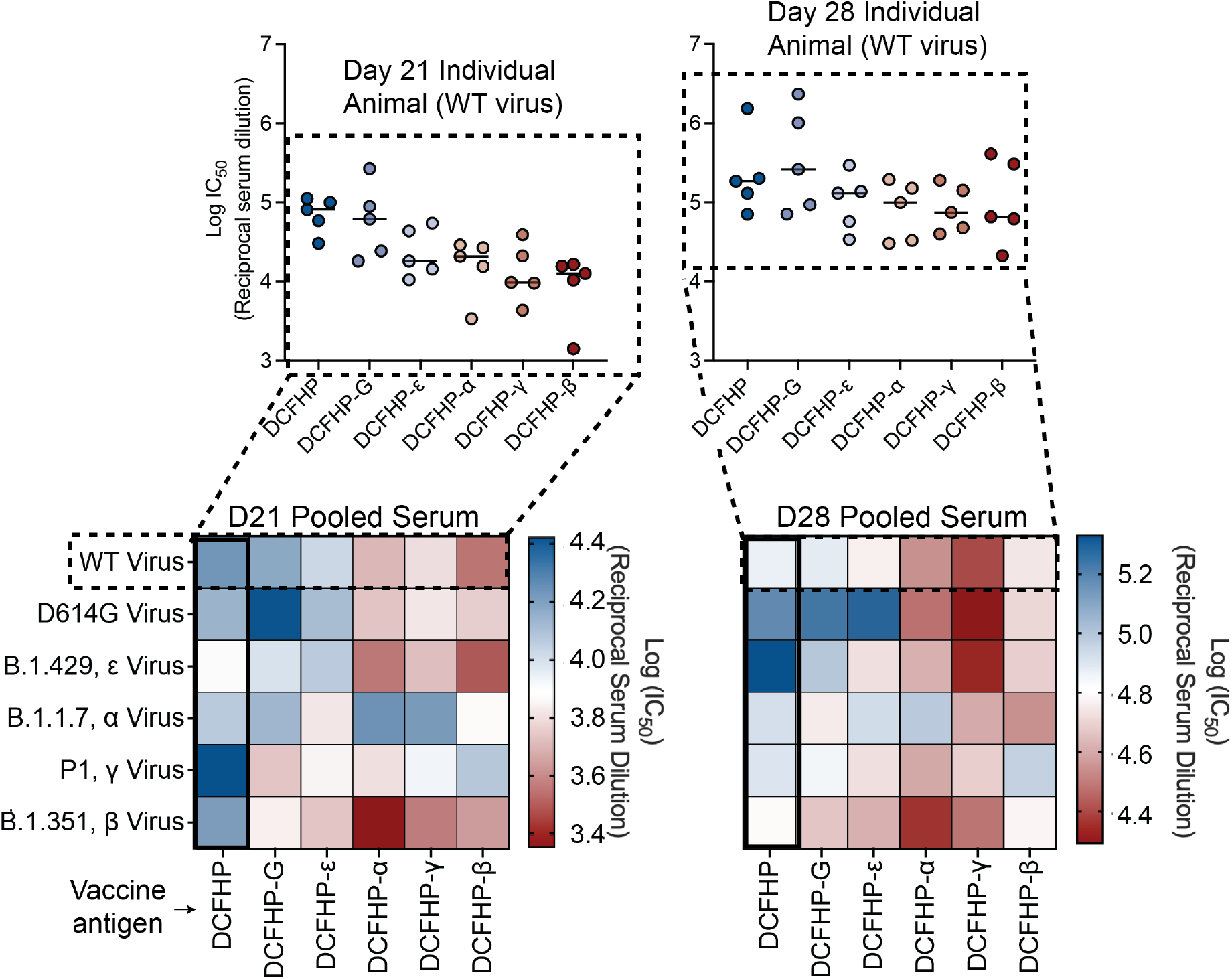
The parent DCFHP vaccine yields the most robust and most cross-reactive neutralizing immune response against VOCs. Pooled sera from mice in each vaccinated group (labeled, bottom), at equal volume ratios from each of n = 5 mice, were tested for neutralization against variant pseudoviruses (labeled, left). Heatmaps showing pooled sera neutralization at day 21 (left) or day 28 (post-boost, right) were generated as for Figure 6, where blue colors indicate more robust neutralization. DCFHP immunized animals elicit the most cross-reactive response. Individual animal neutralization data against the Wuhan-1 sequence are shown in insets at the top and closely resemble the data from the pooled sera analyses in the heatmaps.

Finally, we confirmed these results with a live viral neutralization assay, where the results are qualitatively consistent with those observed with the pseudovirus assay. Notably, we again observed the superiority of the DCFHP vaccine over the DCFHP-variants for neutralizing all VOCs, except for the Epsilon-variant, where the two were comparable (Figure 8). Taken together, these results indicate that the original DCFHP vaccine candidate is the most optimal vaccine antigen, producing neutralizing activity against all of the tested VOCs to a greater extent than the variant-specific vaccines.

**Figure 8.**
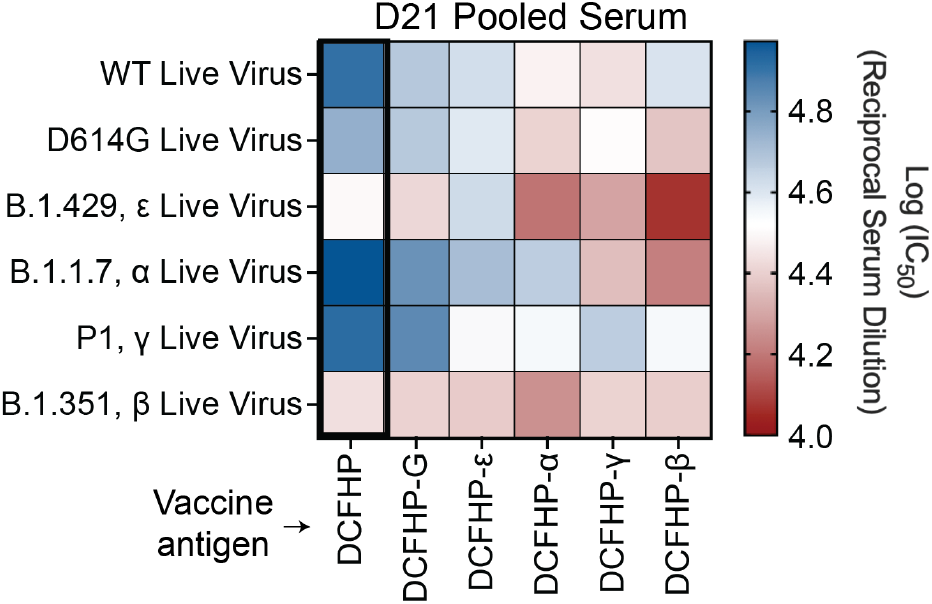
The parent DCFHP produces the most potent neutralizing immune response against live SARS-CoV-2 viruses. Live viral neutralization data were plotted as in Figures 6 and 7, with the pooled serum from vaccination on the x-axis and the live virus on the y-axis, and where blue tones indicate more robust neutralization. The heatmap resembles those in Figure 7 and is consistent with the conclusion that the original parent DCFHP is superior to the DCFHP-variants with respect to eliciting potent, broadly neutralizing antisera.

## DISCUSSION

Our earlier studies demonstrated that DCFHP, a spike-functionalized ferritin-based nanoparticle, forms a stable, homogeneous species with an observed molecular weight of 3.4 MDa^1^. We found here that introduction of amino acid substitutions found in the early SARS-CoV-2 VOCs into the spike component of DCFHP led to an altered nanoparticle homogeneity, particularly for DCFHP-β and DCFHP-γ (Figure 3). This is striking given the number of mutations is rather low relative to the size of the spike protein (less than 1% at most). In addition, the thermal melting transitions of the DCFHP-variants were different from those observed with DCFHP (Figure 5), suggesting that protein stability is impacted. Most strikingly, the ability of DCFHP-variants to elicit neutralizing antisera in mice was generally inferior to that of DCFHP – measured not only against the WT virus, but also against the corresponding VOCs (Figures 7, 8).

Taken together, our results indicate that, at least in the background of DCFHP, predicting the effect of mutations on the immunogenicity of vaccine variants is not as straightforward as what one would anticipate for introducing amino acid substitutions into a rigid protein. Rather, it appears likely that these mutations are impacting the structural integrity of the spike protein and/or exposing non-neutralizing epitopes, with consequences for immunogenicity. This result is particularly striking given that these immunizations were done with purified, homogenous nanoparticles – utilizing a platform amenable to protein purification, unlike some other vaccine platforms. Earlier studies have reported that the SARS-CoV-2 spike is dynamic, and not a rigid protein^23-26^. Taken together, these results suggest that, for the creation of optimal vaccines against VOCs, it may be important to profile each mutation individually for their impact on protein structure and immunogenicity.

An outstanding question is whether these results have implications for mRNA vaccines based on VOCs. There are important caveats to state from the outset. Our results: (i) were generated with DCFHP, a protein nanoparticle-based vaccine, (ii) were obtained in mice, (iii) studied only VOCs and DCFHP-variants that occurred before the Delta and Omicron VOCs, and (iv) utilized the background of the stabilizing HexaPro mutations^2^, instead of the 2P version of spike^11,12^ found in approved mRNA vaccines. Nonetheless, the striking conclusion from our studies is that of the DCFHP-variants tested, the original DCFHP vaccine, based on the Wuhan-1 sequence, was the optimal construct for eliciting neutralizing antisera against all VOCs tested. Biochemical and biophysical characterization of the DCFHP-variants reveal differences from DCFHP that may be responsible for these immunological differences. We note that in the case of mRNA vaccines, it has not been possible to our knowledge to characterize the structure, homogeneity, and stability of the expressed proteins. Existing evidence indicates that, surprisingly, variant-specific mRNA vaccines have similar or only slightly better efficacy against the VOCs that they were based upon, as compared to vaccines based on the original Wuhan-1 sequence^27-31^. These observations, taken together with the results presented here, suggest that caution might be appropriate before assuming that variant-chasing strategies are necessarily the best for protection against future VOCs. Finally, our results may have important implications for the choice of a vaccine to establish optimal immune imprinting^32,33^ (e.g., in infants) to protect against sarbecoviruses, as variant-specific vaccines may result in narrowing of the immune response.

## ACKNOWLEDGEMENTS

We thank L. Borio, M. Bucci, M. Kay, S. Kim, P. Krause, B. Palanski, A. Powell, D. Xu, A. Utz, for helpful comments on an earlier draft of this manuscript. The findings and conclusions in this article are those of the author(s) and do not necessarily represent the views or opinions of the California Department of Public Health or the California Health and Human Services Agency. This work was supported by the Frank Quattrone & Denise Foderaro Family Research Fund, the Chan Zuckerberg Biohub, the Stanford Innovative Medicines Accelerator, the Virginia & D.K. Ludwig Fund for Cancer Research, and an NIH Director’s Pioneer Award (DP1AI158125) to P.S.K.

## COMPETING INTERESTS

P.A.B.W., M.S., N.F., and P.S.K. are named as inventors on patent applications applied for by Stanford University and the Chan Zuckerberg Biohub on immunogenic coronavirus fusion proteins and related methods, which have been licensed to Vaccine Company, Inc. P.A.B.W. is an employee of, and P.S.K. is a co-founder and member of the Board of Directors of Vaccine Company, Inc. All other authors declare no competing interests.

## Materials and Methods

### IgG plasmids

Antibody sequences and Fc-tagged ACE2 were cloned into the CMV/R plasmid backbone for expression under a CMV promoter. The antibodies with variable HC/LC were cloned between the CMV promoter and the bGH poly(A) signal sequence of the CMV/R plasmid to facilitate improved protein expression. The variable region was cloned into the human IgG1 backbone. This vector also contained the HVM06_Mouse (P01750) Ig HC V region 102 signal peptide to allow for protein secretion and purification from the supernatant.

### Lentivirus plasmids

The 21-amino acid C-terminally truncated spike proteins with native signal peptides were cloned in place of the HDM-SARS2-spike-delta21 gene (Addgene plasmid, 155130). This construct contains a 21-amino acid C-terminal deletion to promote viral production, contained in all SARS-CoV-2 variants of concern. The other viral plasmids that were used have been previously described^34^, including pHAGE-Luc2-IRES-ZsGreen (NR-52516), HDM-Hgpm2 (NR-52517), pRC-CMV-Rev1b (NR-52519) and HDM-tat1b (NR-52518).

### Other plasmids

An in-house pADD2 vector was used for all nanoparticle production. Sequences encoding DCFHP (residues 1–1146 of HexaPro) followed by an SGG linker and then the ferritin protein^1^ were cloned into the pADD2 vector backbone using HiFi PCR (Takara) followed by In-Fusion (Takara) cloning with EcoRI/XhoI cut sites. This was followed by an amplicon containing H. pylori ferritin (residues 5–168) originally generated as a gene-block fragment from Integrated DNA Technologies (IDT). VOCs mutations were cloned into the DCFHP vector by stitching PCR. Briefly, overlapping fragments encoding mutations were produced by developing forward and reverse primers encoding the mutation(s) and amplifying gene fragments with the whole gene reverse and a whole gene forward primer respectively – if multiple mutations were being introduced concurrently, adjacent mutation reverse/forward primers were utilized. These resulting fragments were then gel extracted and “stitched” by overlap extension PCR for 5 cycles, followed by amplification with the whole gene forward and whole gene reverse primers. Final gene products were gel extracted and cloned using In-Fusion. All PCR was done with HiFi PCR master mix (Takara) following the manufacturer recommendations.

### Protein production

Transiently expressed proteins were expressed in Expi293F cells. Expi293F cells were cultured in media containing 66% Freestyle/33% Expi media (Thermo Fisher Scientific) and grown in TriForest polycarbonate shaking flasks at 37 °C in 8% CO2. The day before transfection, cells were resuspended to a density of 3 × 10^6^ cells/mL in fresh media. The next day, cells were diluted and transfected at a density of approximately 3–4 × 10^6^ cells/mL. Transfection mixtures were made by combining the following components and adding the mixture to cells: maxi-prepped DNA, culture media, and FectoPro (Polyplus) at a ratio of 0.5–0.8 µg:100 µl:1.3 µl to 900 µL cells. For example, for a 100-ml transfection, 50–80 µg of DNA would be added to 10 mL of culture media, and then 130 µl of FectoPro would be added to this. After mixing and a 10-minute incubation, the resultant transfection cocktail would be added to, for example, 90 mL of cells. The cells were harvested 3–5 days after transfection by spinning the cultures at >7000 × *g* for 15 min. Supernatants were filtered using a 0.22-µm filter.

### Protein purification—Fc Tag-containing proteins

All proteins containing an Fc tag (for example, IgGs and hFc-ACE2) were purified using a 5-ml MabSelect SuRe PRISM column on the ÄKTA pure fast protein liquid chromatography (FPLC) system (Cytiva). Filtered cell supernatants were diluted with a 1/10 volume of 10× PBS. The ÄKTA system was equilibrated with: A1—1× PBS; A2—100 mM glycine pH 2.8; B1—0.5 M NaOH; Buffer line—1× PBS; and Sample lines—H_2_O. The protocol washes the column with A1, followed by loading of the sample in Sample line 1 until air is detected in the air sensor of the sample pumps, followed by 5 column volume washes with A1 and elution of the sample by flowing of 20 ml of A2 (directly into a 50-ml conical containing 2 ml of 1 M Tris pH 8.0). The column was then regenerated with 5 column volumes of A1, B1, and A1. The resultant Fc-containing samples were concentrated using 50-kDa or 100-kDa cutoff centrifugal concentrators.

### Nanoparticle purification

DCFHP purification was done first via flowthrough anion exchange followed by size-exclusion. Two buffers were initially prepared (buffer A: 20 mM Tris pH 8.0, buffer B: 20 mM Tris pH 8.0, 1 M NaCl). Filtered Expi293F or CHO supernatant was diluted with buffer B by 1/5 volume to a final concentration of 200 mM NaCl. The HiTrap® Q anion exchange column was washed with 5 column volumes (CV) sequentially with buffers A, B, A, prior to sample loading. Diluted sample containing 200 mM NaCl was added to the column and the flow through was collected. One 5 mL HiTrap® Q anion exchange column was used for every 200 mL of diluted media. Multiple columns were joined in series for larger sample volumes. 100 kDa MWCO Amicon® spin filters were used to concentrate and buffer exchange the sample with 2 washes with 20 mM Tris, 150 mM NaCl pH 7.5. After the final wash, the sample was concentrated and filtered with a 0.22 µm filtered. The filtered sample was then loaded onto an SRT SEC-1000 column pre-equilibrated with 20 mM tris pH 7.5, 150 mM NaCl. The fractions from the beginning of the nanoparticle peak were pooled. Samples were routinely concentrated to 0.5–1 mg/mL and flash frozen in 20 mM Tris, 150 mM NaCl, 5% sucrose (weight:volume) buffer.

### SDS-PAGE analysis

For the SDS-PAGE quantitation of nanoparticles, expression media samples were diluted with 2X reducing SDS sample buffer at 1:1 ratio and the mixture was loaded on a 10-well 4–20% SDS-PAGE gradient gel (BioRad). Image J software was used to quantify the signal compared to the negative control (mock transfected culture). The band intensity for each nanoparticle was normalized to the WT DCFHP signal.

### SEC analysis of ferritin-antigen nanoparticles

SEC was performed on an Agilent 1260 Infinity II HPLC instrument. The purified antigen (20 μg) was loaded onto a SRT SEC-1000 4.6 mm × 300 mm column and equilibrated in 1× PBS (pH 7.4). SRT SEC-1000 column was run at a flow rate of 0.35 mL/min. Data were plotted using Prism Version 9.4.1.

### Biolayer interferometry (BLI) of mAbs binding to SARS-CoV-2 purified antigens

BLI was performed on an OctetRed 96 system (ForteBio). Samples were assayed in “Octet buffer” (0.5% bovine serum albumin, 0.02% Tween, 1×DPBS (1xDPBS from Gibco)) in 96-well flat-bottom black-wall, black-bottom plates (Greiner). Biosensors were equilibrated in Octet buffer for at least 10 min and regenerated in 100 mM glycine (pH 1.5) prior to sample testing. Tips in experiments that involved regeneration were regenerated in 100 mM glycine (pH 1.5) prior to testing. Anti-Human Fc sensor tips (ForteBio, now Sartorius) were loaded with 200 nM mAb or FcACE2 to a threshold of 0.8 nm and then submerged into wells containing 100 nM (protomer/monomer concentration) of each antigen. Data were plotted using Prism Version 9.4.1.

### Differential scanning fluorimetry

Thermal melts were determined using the Prometheus NT.48 (Nanotemper). Samples were loaded into Prometheus NT.Plex nanoDSF Grade High Sensitivity Capillary Chips and the laser intensity was set such that the discovery scan placed the auto fluorescence between the upper and lower bounds. Samples were allowed to melt using the standard melt program (1 °C/min). Melting temperatures were determined by peaks on the first derivatives of the ratio of F350/F330. Data were plotted using Prism Version 9.4.1.

### Mouse immunizations

Balb/c female mice (6–8 weeks old) were purchased from The Jackson Laboratory. All mice were maintained at Stanford University according to the Public Health Service Policy for “Humane Care and Use of Laboratory Animals” following a protocol approved by Stanford University Administrative Panel on Laboratory Animal Care (APLAC-33709). Mice were immunized intramuscularly with 5μg DCFHP or DCFHP-variant at days 0 and 21 with alum/CpG (which contains 500 μg alum + 20 μg CpG). All antigen doses were formulated in Tris Buffer (20 mM, pH 7.5, 150 mM NaCl, 5% sucrose) in a total volume of 100 μL per injection. DCFHP and DCFHP-variants were produced in Expi293F cells via transient transfection for all mouse experiments. Serum was collected and processed using Sarstedt serum collection tubes. Mouse serum was centrifuged at 10,000 × g for 15 min and heat inactivated for 30 min at 56 °C. For pooled serum samples, equi-volume mixtures of antisera was produced and subsequently heat inactivated.

### Lentivirus production

SARS-CoV-2 VOCs and SARS-CoV-1 spike pseudotyped lentiviral particles were produced. HEK293T cells were transfected with plasmids described above for pseudoviral production using BioT transfection reagent. Six million cells were seeded in D10 media (DMEM + additives: 10% FBS, L-glutamate, penicillin, streptomycin, and 10 mM HEPES) in 10-cm plates 1 day before transfection. A five-plasmid system (plasmids described above) was used for viral production, as described in Crawford et al^34^. The spike vector contained the 21-amino acid truncated form of the SARS-CoV-2 spike sequence from the Wuhan-Hu-1 strain of SARS-CoV-2 or VOCs. VOCs were based on wild-type (WT) (Uniprot ID: BCN86353.1); Alpha (sequence ID: QXN08428.1); Beta (sequence ID: QUT64557.1); Gamma (sequence ID: QTN71704.1), mutations shown in Fig. 1. The plasmids were added to D10 medium in the following ratios: 10 µg pHAGE-Luc2-IRS-ZsGreen, 3.4 µg FL spike, 2.2 µg HDM-Hgpm2, 2.2 µg HDM-Tat1b and 2.2 µg pRC-CMV-Rev1b in a final volume of 1000 µl. To form transfection complexes, 30 µl of BioT (BioLand) was added. Transfection reactions were incubated for 10 min at room temperature, and 9 ml of medium was added slowly. The resultant 10 ml was added to plated HEK cells from which the medium had been removed. Culture medium was removed 24 h after transfection and replaced with fresh D10 medium. Viral supernatants were harvested 72 h after transfection by spinning at 300 × *g* for 5 min, followed by filtering through a 0.45-µm filter. Viral stocks were aliquoted and stored at −80 °C until further use.

### Neutralization

The target cells used for infection in viral neutralization assays were from a HeLa cell line stably overexpressing the SARS-CoV-2 receptor, ACE2, as well as the protease known to process SARS-CoV-2, TMPRSS2. Production of this cell line is described in detail in ref.^35^ with the addition of stable TMPRSS2 incorporation.

ACE2/TMPRSS2/HeLa cells were plated 1 day before infection at 10,000 cells per well. Ninety-six-well white-walled, clear-bottom plates were used for the assay (Thermo Fisher Scientific) and a white seal was placed on the bottom prior to readout. As previously described^7^, on the day of the assay, dilutions of heat inactivated serum were made into sterile D10 medium. Samples were analyzed in technical duplicate in each experiment. Virus-only wells and cell-only wells were included in each assay. For pooled serum samples, equi-volume mixtures of antisera was produced and heat inactivated. The resultant sera was then serially diluted in media and used in neutralization assays.

A virus dilution was made containing the virus of interest (for example, SARS-CoV-2) and D10 media (DMEM + additives: 10% FBS, L-glutamate, penicillin, streptomycin, and 10 mM HEPES). Virus dilutions into media were selected such that a suitable signal would be obtained in the virus-only wells. Polybrene was present at a final concentration of 5 µg/mL in all samples. 60 µL of heat inactivated sera was mixed with 60 µL viral dilution to make a final volume of 120 µl in each well.

The inhibitor (serum dilution) and virus mixture was left to incubate for 1 h at 37 °C. After incubation, the medium was removed from the cells on the plates made 1 day prior. This was replaced with 100 µl of inhibitor/virus dilutions and incubated at 37 °C for approximately 48 h. Infectivity readout was performed by measuring luciferase levels.

48 h after infection, half the medium was removed from all wells and cells were lysed by the addition of 50 µl of BriteLite assay readout solution (Perkin Elmer) to make a a 1:1 dilution into each well. Luminescence values were measured using a BioTek Synergy HT (BioTek) or Tecan M200 microplate reader. Each plate was normalized by averaging cell-only (0% infectivity) and virus-only (100% infectivity) wells. Cell-only and virus-only wells were averaged. Normalized values were fit with a four-parameter non-linear regression inhibitor curve in Prism to obtain 50% neutralizing titer (NT_50_) values.

### ELISA

Recombinant trimeric spike proteins (5 µg/mL) were plated in 50 µl in each well on a MaxiSorp (Thermo Fisher Scientific) microtiter plate in 1xPBS and left to incubate for at least 1 h at room temperature. These were washed 3 times with 300 µl of ddH_2_O using an ELx 405 Bio-Tex plate washer and blocked with 150 µl of ChonBlock (Chondrex) for at least 1 h at room temperature. Plates were washed 3x with 300 µl of 1x PBST. Mouse serum samples, serially diluted in diluent buffer (1x PBS, 0.1% Tween) starting at 1:50 serum dilution was then added to coated plates for 1 h at room temperature. This was then washed 3x with PBST. HRP goat anti-mouse (BioLegend 405306) was added at a 1:10,000 dilution in diluent buffer for 1 h at room temperature. This was left to incubate at room temperature for 1 h and then washed 6x with PBST. Finally, the plate was developed using 50 µl of 1-Step TMB Turbo-TMB-ELISA Substrate Solution (Thermo Fisher Scientific) per well, and the plates were quenched with 50 µl of 2 M H_2_SO_4_ to each well. Plates were read at 450 nm and normalized for path length using a BioTek Synergy HT Microplate Reader.

### Live SARS-CoV-2 virus isolation and passages

Variants were obtained from two sources, the WRCEVA collection (WA-1/2020) or from de-identified nasopharyngeal (NP) swabs sent to the California Department of Public Health from hospitals in California for surveillance purposes. To isolate from patient swabs, 200 µl of an NP swab sample from a COVID-19-positive patient that was previously sequence-identified was diluted 1:3 in PBS supplemented with 0.75% BSA (BSA-PBS) and added to confluent Vero E6-TMPRSS2-T2A-ACE2 cells in a T25 flask, allowed to adsorb for 1 h, inoculum removed, and additional media was added. The flask was incubated at 37 °C with 5% CO_2_ for 3–4 days with daily monitoring for cytopathic effects (CPE). When 50% CPE was detected, the contents were collected, clarified by centrifugation and stored at −80 °C as a passage 0 stock. 1:10 diluted passage 0 stock was used to inoculate Vero E6-TMPRSS2-T2A-ACE2 grown to confluency in T150 flasks and harvested at approximately 80% CPE. All viral stocks were sequenced to confirm lineage, and 50% tissue culture infectious dose (TCID_50_) was determined by titration.

### Live SARS-CoV-2 virus 50% CPE endpoint neutralization

CPE endpoint neutralization assays were performed following the limiting dilution model using sequence-verified viral stocks of WA-1, and VOCs in Vero E6-TMPRSS2-T2A-ACE2. Three-fold serial dilutions of inhibitor (antisera) were made in BSA-PBS and mixed at a 1:1 ratio with 100 TCID_50_ of each virus and incubated for 1 h at 37 °C. Final inhibitor dilutions ranged from 500 nM to 0.223 nM. Then, 100 µl of the plasma/virus mixtures were added in duplicate to flat-bottom 96-well plates seeded with Vero E6-TMPRSS2-T2A-ACE2 at a density of 2.5 × 10^4^ per well and incubated in a 37 °C incubator with 5% CO_2_ until consistent CPE was seen in the virus control (no inhibitor added) wells. Positive and negative controls were included as well as cell control wells and a viral back titration to verify TCID_50_ viral input. Individual wells were scored for CPE as having a binary outcome of ‘infection’ or ‘no infection’, and the IC_50_ was calculated using the Spearman–Karber method. All steps were done in a Biosafety Level 3 laboratory using approved protocols.

## Notes

### Summary of Updates

The top labels on Figure 7 were incorrect, they have been corrected in this version.

